# Linkage of a plasma zinc signature and impaired insulin receptor activation: Implications for the mechanism of type 2 diabetes mellitus

**DOI:** 10.1101/849091

**Authors:** Shashidhar M. Prabhakar

## Abstract

Type 2 diabetes mellitus (T2DM) is characterized by decreased plasma zinc levels and hyperzincuria, yet the underlying cause of these zinc disturbances is unknown. In this study, we compared postprandial plasma zinc levels in samples from T2DM and healthy control subjects to determine whether zinc is associated with a different set of proteins. We found that in T2DM a considerable amount of zinc remained following albumin/Ig depletion. A discrepancy in total amount of zinc in the remaining set of zinc-associated proteins identified and estimated by protein analysis as alpha-2 macroglobulin (A2M), and in T2DM alone some bacterial proteinases as well, indicated that the likely source of this discrepancy was from bacterial zinc proteinases trapped by A2M that obscured the high levels of these proteinases. Furthermore, an insulin receptor assay examined whether activated A2M (A2MFF) affected insulin receptor activation. The results showed a significant decrease in insulin receptor activation following repeated treatments with A2MFF but not after a single treatment with A2MFF. Our findings suggest that in T2DM, A2MFF likely arises from the trapping of zinc-dependent bacterial proteinases and impairs insulin receptor activation from a prolonged presence, which may result in a “receptor-protective” effect manifested as insulin resistance.

## Introduction

Type 2 diabetes mellitus (T2DM) is increasing worldwide due, in part, to rising rates of obesity caused by poor diet [1] and physical inactivity. The global prevalence of diabetes among adults over 18 years of age has increased from 4.7% in 1980 to 8.5% (422 million) in 2014 [2].

While drugs for the management of T2DM are numerous, the lack of a thorough understanding of the underlying causes of T2DM has prevented the development of a cure for the condition. Numerous studies in the last several decades have observed decreased plasma zinc levels [3], loss of zinc [4] and effects of zinc supplementation on glycemic control [5] in T2DM, but the underlying causes of the disturbances in zinc homeostasis or the role in T2DM remains unclear.

The aim of this study was to identify the source of decreased zinc and zinc loss and to discover a mechanism for the loss of zinc observed in T2DM. This study compared postprandial plasma zinc levels between healthy controls and T2DM subjects to determine possible differences in zinc-associated proteins. This study has identified the likely source of zinc loss as bacterial zinc proteinases that cause the formation of activated A2M, which is then presumably cleared through the LRP1 receptor pathway. Additionally, we also found that in the continued presence of activated A2M, activation of the insulin receptor is impaired and results in a reduced insulin response.

## Methods and Materials

### Research Design

This study was approved by the New England Institutional Review Board (NEIRB, Needham, MA US). This study recruited and gathered data from 9 volunteers, who included 6 diabetic or prediabetic patients and 3 controls. The panel included one female and one Caucasian, and all subjects were males of South Asian origin. The T2DM volunteer subjects had diabetic or prediabetic ranges of fasting glucose levels above 110 mg/dL, and the control subjects were at 100 mg/dL or below. Subjects were informed of the research procedure and risks of blood draw, and consent was obtained as per NEIRB guidelines. As no interventions were planned or performed, this study was not classified as a clinical trial under the NIH definition. Blood draws and tests were contracted and performed between June 2018 and June 2019 at a nationally recognized certified commercial laboratory facility by professional personnel. Fasting blood draw was performed after overnight fasting, and postprandial blood draw was performed 1 hour after a light breakfast.

### Zinc Test

Zinc level measurement in plasma was determined using a standard clinically approved test number 945, offered by the national commercial laboratory. Blood was drawn at the testing facility following the collection protocol for the zinc test. Both samples of unprocessed and plasma depleted of albumin/Ig at the researchers’ laboratory were returned to the clinical laboratory for testing within the period recommended by guidelines for the zinc test. For zinc level determination, 300 µl of plasma were typically albumin/Ig depleted and then reconstituted with 300 µl of unprocessed plasma into a final volume of 900 µl using 1x phosphate-buffered saline (PBS), resulting in a 3-fold dilution. This sample reconstitution was performed to bring zinc levels above the detection threshold of 10 µg/dL for the liquid chromatography/Mass Spectroscopy (LC/MS) protocol used for zinc level calculation at the commercial clinical laboratory. In addition, two samples of 900 µl, one an unprocessed sample diluted 3x using 1x PBS, and the other an unprocessed, undiluted plasma sample were tested for reference. The results are reported in mg/dL.

### Albumin/Ig Depletion

Plasma obtained from blood collected at fasting and one hour postprandial draw were depleted of albumin/Ig using the PureProteome™ kit (Catalogue Number: LSKMAGHD), purchased from EMD Millipore (Temecula, CA US), according to the manufacturer’s instructions.

### Zinc Immobilized Metal Affinity Chromatography

Zinc Immobilized Metal Affinity Chromatography (Zn-IMAC) columns of 2 ml volume were purchased from Genelinx International Inc/bioWorld (Dublin OH). Albumin/Ig-depleted postprandial plasma from T2DM subjects and healthy controls was filtered, and zinc binding proteins were eluted according to the supplied protocol using HIS-Select wash and elution buffers purchased from Sigma-Aldrich (Saint Louis, MO).

### Liquid Chromatography with Tandem Mass Spectrometry Protein Identification

Poochon Scientific Inc (Frederick, Maryland, USA) performed LC/MS/MS analysis on the proteins with zinc affinity from albumin/Ig depleted plasma, purified by Zn-IMAC, using a Thermo Scientific Q-Exactive hybrid Quadrupole-Orbitrap Mass Spectrometer and a Thermo Dionex UltiMate 3000 RSLCnano System. Samples were shipped at 4c overnight to the facility where samples were stored at −20 until analysis was performed.

### Insulin Response Assay

The PathHunter^®^ Insulin Bioassay Kit was purchased from Discoverx/Eurofins (Fremont, CA, US). This kit was used to measure the insulin response through chemiluminescent output in a cell-based functional assay for insulin receptor isoform b. The insulin response was measured for A2M at a concentration of 1.2 µg/ml with either A2MFF added once, at a concentration of 0.24 µg/µl, or A2MFF added every 30 minutes, at a concentration of 0.24 µg/µl, throughout the 3-hour incubation after the addition of insulin. Native A2M was purchased from Genway Biotech (San Diego, CA US), and A2MFF was purchased from Sigma-Aldrich (St. Louis, MO US).

### Statistical Analysis

Statistical differences in plasma zinc levels between experimental groups (Fig 1) were determined by paired, two-tailed Student’s *t-*test, and values of *P* < 0.05 were considered significant. Statistical differences in the insulin receptor activation in response to insulin (Fig 2) were assessed by paired, two-tailed Student’s *t-*test two-way ANOVA followed by Dunnett’s multiple comparisons test. In both methods values of *P* < 0.05 were considered significant. Statistical analysis was performed using GraphPad Prism version 8.2.1 for MacOS, GraphPad Software (San Diego, CA US), www.graphpad.com.

**Fig 1.**
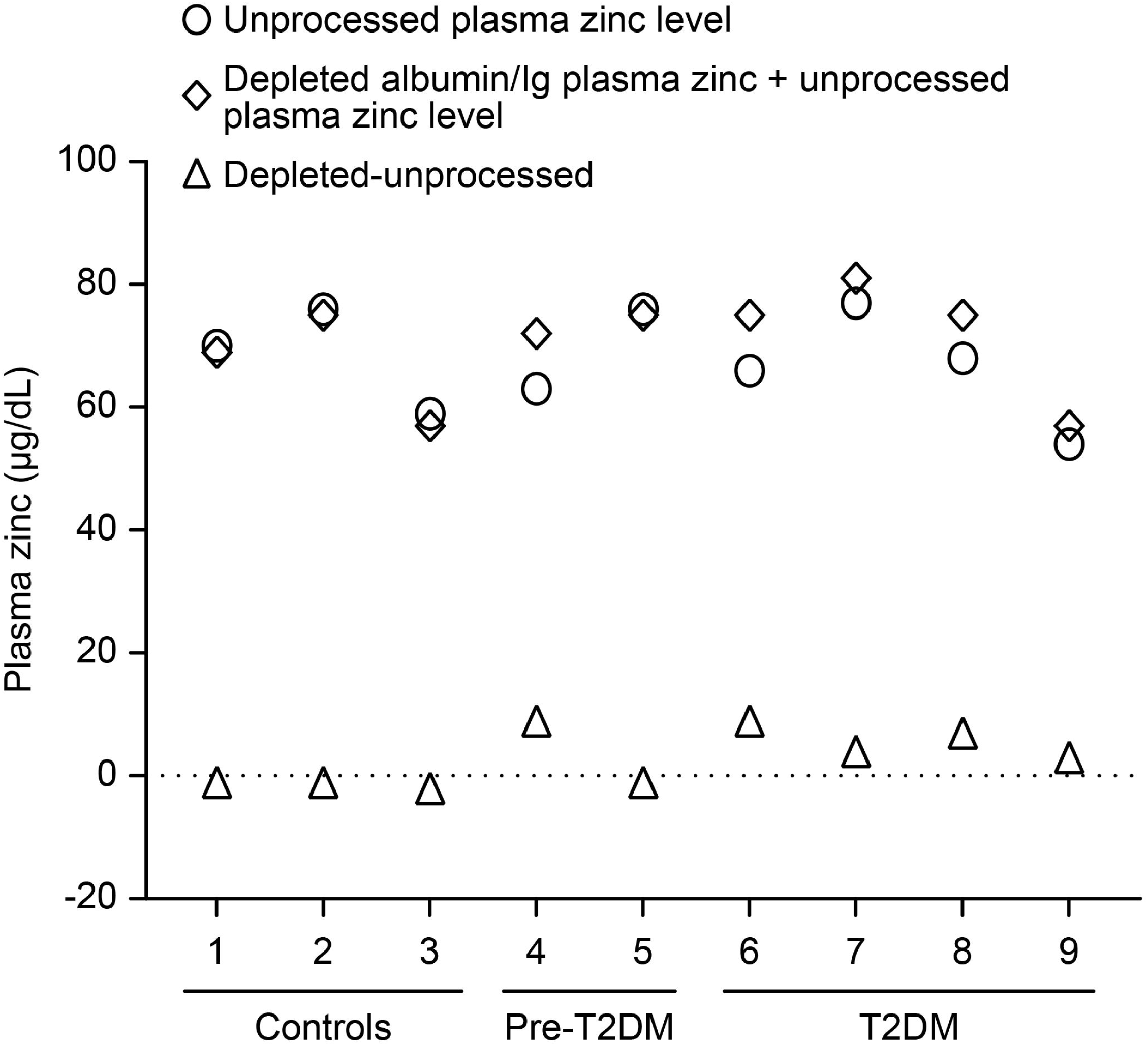
Scatter plot of the zinc levels from unprocessed plasma (predepletion) and albumin/Ig-depleted plasma (postdepletion) in the controls and T2DM subjects. The difference in zinc levels is indicated by open levels. Samples 1–3 are the controls, and samples 4–9 are the T2DM samples (including prediabetes). There was no significant difference between the predepletion and postdepleted plasma in the controls (paired two-tailed *t*-test, *P* = 0.0572), while the T2DM samples showed a significant difference in zinc levels between predepletion and postdepletion plasma (paired two-tailed *t*-test, *P* = 0.0232

**Fig 2.**
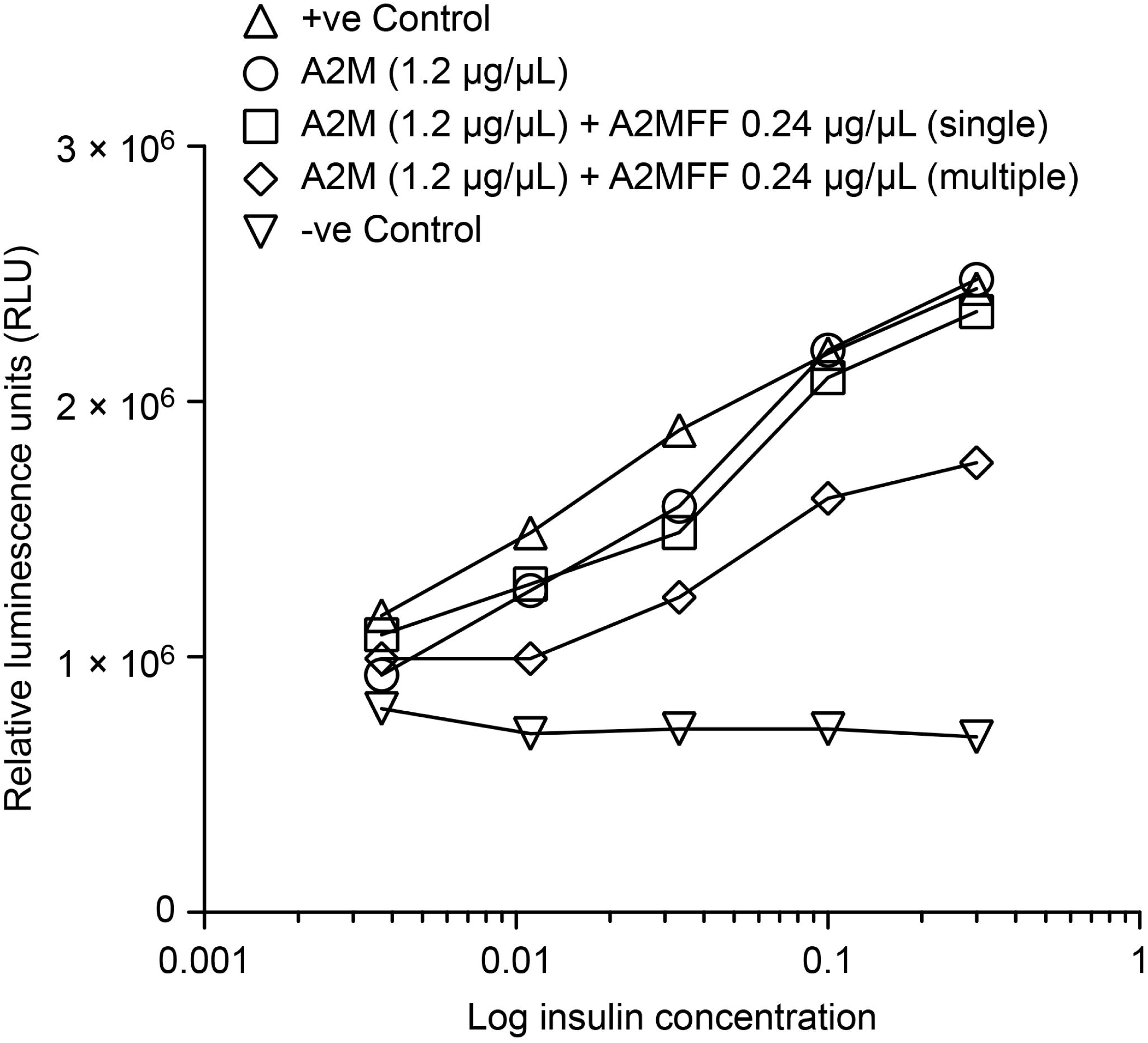
Insulin receptor activation by insulin was measured for A2M and activated, fast form A2M (A2MFF). A2MFF was either added once (single treatment) or added every 30 minutes during a 3-hour incubation (multiple treatments). There was a significant difference between single treatment with A2MFF and multiple treatments with A2MFF in the presence of A2M. A single treatment with A2MFF showed no significant difference when compared to treatment with A2M alone (two-way ANOVA, *P* = 0.9548, *P* = 0.9549). There is a significant difference in insulin receptor activation with multiple treatments with A2MFF when compared to treatment with A2M alone (paired two-tailed *t-*test *P* = 0.0173, two-way ANOVA, *P* = 0.0027, *P* = 0.0148).

## Results

### Comparison of Postprandial Zinc Levels

We measured postprandial plasma zinc levels in subjects with T2DM and healthy controls before and after albumin/Ig depletion. We found zinc levels in all subjects before albumin/Ig depletion were in the normal range of 60–130 µg/dL. Zinc levels after albumin/Ig depletion were nearly zero in the controls and significantly above zero in the T2DM subjects. This is shown as the difference between zinc levels between unprocessed plasma and depleted plasma with a known quantity of added plasma to bring values above the detection threshold (Fig 1). In the controls (samples 1–3), there was no significant difference between pre- and postdepletion plasma (paired two-tailed *t*-test, *P* = 0.0572,). The T2DM group (samples 4–9), however, showed a significant difference in pre- and postdepletion zinc levels (paired two tailed *t*-test, *P* = 0.0232). A comparison of the controls vs. T2DM groups showed a significant difference between the groups of 6.5 units (paired two-tailed *t-*test, *P* = 0.0280). A Gaussian distribution was assumed even though the sample sizes were small. These findings suggest that there is a small fraction of proteins with zinc affinity other than albumin/Ig in T2DM that are not present in healthy controls.

### Liquid Chromatography with Tandem Mass Spectrometry Based Protein Identification

To identify the zinc-associated proteins in the samples after albumin/Ig depletion, we analyzed postprandial albumin/Ig-depleted plasma from each subject after filtration using Zn-IMAC and liquid chromatography with tandem mass spectrometry (LC/MS/MS) for protein identification and analysis. Of the 7 samples analyzed, 3 control subjects and 2 prediabetic patient showed zero percent of bacterial proteinases, while 2 T2DM samples showed a minor amount of bacterial proteinases. In the T2DM samples, the bacterial proteinases alone did not add up to the amount of remaining zinc not associated with albumin/Ig. All samples showed that the majority protein was alpha 2 macroglobulin (A2M) (Table 1). While A2M is increased in diabetes, it has a decreased trypsin binding capacity [6], and thus, the most likely source for the remaining zinc levels is from A2MFF by zinc bacterial proteinases trapped by A2M, which is a pan-protease inhibitor. Human and bacterial sequences were searched and identified using Proteome Discoverer 2.2 software.

**Table 1.** Relative % of Alpha 2 Macroglobulin and Bacterial Proteases Detected (Excluding A. lyticus). Only T2DM Subjects Showed some Zinc Bacterial Proteases.

| Relative % of zinc | Control -1 | Control -2 | Control -3 | T2DM-1 | T2DM-2 (Pre-diabetic) | T2DM-3 (Pre-diabetic) | T2DM-4 |
| --- | --- | --- | --- | --- | --- | --- | --- |
| alpha-2 macroglobulin | 92.5 | 98.0 | 99.2 | 83.5 | 97.2 | 92.2 | 95.7 |
| Total of next 5 highest levels of proteins | 4.6 | 2.6 | 1.1 | 12.3 | 2.7 | 2.6 | 3.4 |
| Bacterial proteinases (excluding A. lyticus) | 0 | 0 | 0 | 0.06 | 0 | 0 | 0.02 |

### Measurement of Insulin Receptor Activation

The effect of non-activated A2M and activated, fast form A2M (A2MFF) on insulin activity was measured. Commercially available purified A2M and A2MFF were subjected to one of two treatments at varying insulin concentrations. Treatment consistent of a single dose of activated A2M (A2MFF single) or multiple additions of activated A2M (A2MFF multiple) every 30 minutes for the duration of the 3-hour incubation in the presence of 1.2 µg/ml of non-activated A2M, at a concentration of 0.24 µg/µl, which approximately a fifth of the included A2M concentration. Insulin receptor activation was assessed by measuring relative luminescence units using the β-galactosidase assay of the PathHunter^®^ insulin receptor activation assay. The results showed a large, significant decrease in insulin receptor activation following A2MFF treatment every 30 minutes relative to a single treatment of A2MFF (paired two-tailed *t-*test, *P* = 0.0173). There was no significant difference between the effect of A2M alone and A2M with a single A2MFF treatment on insulin receptor activation (paired two-tailed *t-*test, *P* = 0.0615) (Fig 2). We performed a two-way ANOVA analysis and observed a significant difference in insulin receptor activation at two different concentrations (300 ng/mL, 100 ng/mL) between A2M with a single treatment of A2MFF (two-way ANOVA, *P* = 0.9548, *P* = 0.9549), which showed no significant difference compared with A2M alone, and A2M with multiple treatments of A2MFF (two-way ANOVA, *P* = 0.0027, *P* = 0.0148), which showed in a significant difference compared to A2M alone (Fig 2). These results demonstrate that insulin receptor activation is significantly reduced in the presence of A2MFF, likely due to interference with the ability of insulin to access the insulin receptor.

We also performed insulin receptor activation assays individually with A2M, A2MFF. We found there was no significant difference in insulin receptor activation between A2M and a single treatment with A2MFF (paired two-tailed *t*-test, *P* = 0.2514). Furthermore, in a separate experiment, we used almost twice the concentration of A2MFF (0.44 µg/µl) compared to that in a previous experiment, with an increased volume (15% of the reaction volume compared to 3% of the reaction volume in the previous experiment), in the presence of A2M. We found a similar reduction in insulin receptor activation following both single (paired two-tailed *t-*test, *P* = 0.0043) and multiple treatments with A2MFF (paired two-tailed *t-*test, *P* = 0.0016) when compared with A2M alone. The reduction in insulin receptor activation in the large single treatment group was prolonged and in the same range as that of the group with multiple treatments of A2MFF (data not shown).

## Discussion

A prominent characteristic of T2DM is decreased plasma zinc levels and hyperzincuria, for which a cause is unknown [7]. We hypothesized that zinc loss is due to the association of zinc with a different set of proteins that may be in a biologically unavailable form. To explore this possibility and considering that a majority of zinc in plasma is associated with albumin/Ig [7,8], we compared postprandial albumin/Ig-depleted plasma zinc levels between healthy controls and T2DM subjects. Our results showed some amount of remaining zinc in albumin/Ig depleted plasma in T2DM subjects but not in healthy controls (Fig 1). This finding indicated that a small but significant portion of zinc is bound to proteins other than albumin/Ig in T2DM.

To identify the proteins associated with the remaining zinc in albumin/Ig depleted postprandial plasma of T2DM subjects, we purified the depleted plasma on a zinc immobilized metal affinity chromatography (Zn-IMAC) column and analyzed the proteins using LC/MS/MS. The results showed that the majority of protein was A2M in both the T2DM subjects and controls. This finding is consistent with similar observations in a previous study [8]. Some bacterial proteinases were observed in 2 T2DM subjects but none in healthy controls or 1 prediabetic subject (Table 1). The total amount of bacterial zinc proteinases did not add up to the level of remaining zinc. Prediabetic sample were not different from the control samples for bacterial zinc proteinases, even though we detected some remnant zinc after albumin/Ig depletion. A previous study showed increased A2M levels with a reduced trypsin binding capacity in diabetes [6]. We inferred that these results, albeit indirectly, indicated that the remaining zinc after albumin/Ig depletion likely came from activated A2M proteins that have trapped bacterial zinc proteinases, since A2M is a pan-protease inhibitor that traps free proteinases, resulting in A2MFF. In the prediabetic case, free bacterial proteinases may not be detected because they may be obscured when captured by A2M, without exceeding the capture threshold for levels of activated A2M [9].

Studies have shown that proteinase-activated A2M (A2MFF) binds numerous receptors, including, LRP1, growth factors and cytokines [10]. We hypothesized that A2MFF may also bind the insulin receptor. We tested this possibility with an insulin receptor activation assay. We measured insulin receptor activation following treatment with non-activated A2M with either single or multiple treatments with activated A2M (A2MFF). Since activated A2M is rapidly cleared away by the LRP1 receptor in approximately 2-4 minutes [11,12], multiple treatments with activated A2M were used to replace activated A2M that would be cleared away during the long incubation, thereby providing a continuous presence of activated A2M and revealing its effects.

We observed a significant decrease in insulin receptor activation after multiple treatments with A2MFF (0.24 µg/µl) compared to a single treatment with A2MFF (0.24 µg/µl) (Fig 2). We also observed a decrease in insulin receptor activation with a single high dose (0.44 µg/µl) of A2MFF (data not shown), similar to the results following multiple treatments with A2MFF at a lower concentration. This result demonstrates that the presence of A2MFF is causing an impairment of insulin receptor activation tested in a dose response to insulin. While A2MFF may interact with insulin itself, since A2MFF has been shown to bind cell surface receptors like the NGS receptor [13], we suggest that A2MFF likely binds the insulin receptor. This action would help protect the insulin receptor from the detrimental actions of free bacterial proteinases. Loss of their insulin receptors would be catastrophic for cells; therefore, this critical “receptor-protective” action by A2MFF would be effective as long as bacterial proteinase-activated A2M and therefore likely destructive free proteinases were present. Then, the “receptor-protective” effect would shield the insulin receptor and presumably prevent the activation of the insulin receptor by insulin. Therefore, insulin signaling and response would be diminished as in the phenomenon of insulin resistance. Our hypothesis is supported by a study showing that functional inactivation of IGF-1 and insulin receptors in skeletal muscle causes T2DM [14]. Additionally, myriad effects, such as excessive secretion of glucagon in pancreatic α cells [15] or reduced insulin-mediated potentiation of glucose-stimulated insulin secretion in insulin resistant humans affected by the role of insulin receptor in glucose homeostasis 16,20] and decreases in beta cell mass [17], can be explained by a “receptor-protective” effect of A2MFF on the insulin receptor to shield it from the destructive effect of free bacterial proteinases. We also suggest that the clearance of the bacterial zinc proteinase-activated A2M through the LRP1 receptor is the cause of the observed effect of hyperzincuria and zinc loss in T2DM. Further, the increased LDL levels observed in T2DM could be due to occupation of the LRP1 via clearance of activated A2M, causing a slowdown in turnover of LDL. Furthermore, duodenal mucosal removal abrogates hyperglycemia [18,19], which provides evidence that bacterial zinc proteinases play a central role in T2DM.

There is currently no explanation for a unified hypothesis that links the defects found in T2DM pathogenesis, including insulin resistance and beta cell dysfunction [20]. By providing evidence for the underlying cause, linking well-known and not fully elucidated phenomena in T2DM—zinc loss and insulin resistance—and describing the correlations to related effects, we offer a unifying explanation and demonstrate a possible mechanism of this disease.

In conclusion we believe that the mucosal barrier in the duodenum is compromised due to goblet cell dysfunction or damage caused by antibiotics [21-23]. This phenomenon allows proteinases produced by discrepant gut bacteria to cross the mucosal layer [23] and be absorbed into the bloodstream. Once in the circulation, these bacterial zinc proteinases are bound by A2M, resulting in activated A2M (A2MFF) [6], whose “receptor-protective” action, as we have described, above causes a diminished insulin response that is perceived as insulin resistance and leads to T2DM.

## Supporting information

Supplemental Data Individual Treatment (Ref: Data not shown)

Supplemental Data Single Large Treatment (Ref: Data not shown)

## Acknowledgments

We thank the volunteers who consented to participate in this research study and staff at the clinical testing facility who enabled the service agreement. We acknowledge the help of kkstats in statistical checking and Poochon Scientific for LC/MS/MS-based protein identification. We acknowledge the help of the Springer-Nature editing service and American Journal Experts in manuscript editing. S.P. performed the experiments, analyzed the data and wrote the manuscript.

## References

1. Carroll IM, Maharshak N. Enteric bacterial proteases in inflammatory bowel disease-pathophysiology and clinical implications. World J Gastroenterol. 2013;19: 7531–7543.

2. World Health Organisation. Key facts world health organization. Geneva, Switzerland: World Health Organisation; 2018.

3. Jansen J, Rosenkranz E, Overbeck S, Warmuth S, Mocchegiani E, Giacconi R, et al. Disturbed zinc homeostasis in diabetic patients by *in vitro* and *in vivo* analysis of insulinomimetic activity of zinc. J Nutr Biochem. 2012;23: 1458–1466.

4. Pidduck HG, Wren PJ, Evans DA. Hyperzincuria of diabetes mellitus and possible genetical implications of this observation. Diabetes. 1970;19: 240–247.

5. Jayawardena R, Ranasinghe P, Galappatthy P, Malkanthi R, Constantine G, Katulanda P. Effects of zinc supplementation on diabetes mellitus: A systematic review and meta-analysis. Diabetol Metab Syndr. 2012;4: 13.

6. Roberts RC, Hall PK, Nikolai TF, McKenzie AK. Reduced trypsin binding capacity of α2-macroglobulin in diabetes. Clin Chim Acta. 1986;154: 85–101.

7. Chausmer AB. Zinc, insulin and diabetes. J Am Coll Nutr. 1998;17: 109–115.

8. Yoshino S, Fujimoto K, Takada T, Kawamura S, Ogawa J, Kamata Y, et al. Molecular form and concentration of serum α2-macroglobulin in diabetes. Sci Rep. 2019;9: 12927.

9. Banks RE, Evans SW, Alexander D, Van Leuven F, Whicher JT, McMahon MJ. Alpha 2 macroglobulin state in acute pancreatitis. Raised values of α2 macroglobulin-protease complexes in severe and mild attacks. Gut. 1991;32: 430–434.

10. Cater JH, Wilson MR, Wyatt AR. Alpha-2-macroglobulin, a hypochlorite-regulated chaperone and immune system modulator. Oxid Med Cell Longev. 2019;2019: 5410657.

11. Gonzalez-Gronow M, Cuchacovich M, Llanos C, Urzua C, Gawdi G, Pizzo SV. Prostate cancer cell proliferation *in vitro* is modulated by antibodies against glucose-regulated protein 78 isolated from patient serum. Cancer Res. 2006;66: 11424–11431.

12. Mikhailenko I, Battey FD, Migliorini M, Ruiz JF, Argraves K, Moayeri M, et al. Recognition of α2-macroglobulin by the low density lipoprotein receptor-related protein requires the cooperation of two ligand binding cluster regions. J Biol Chem. 2001;276: 39484–39491.

13. Koo PH, Qiu WS. Monoamine-activated α2-macroglobulin binds trk receptor and inhibits nerve growth factor-stimulated trk phosphorylation and signal transduction. J Biol Chem. 1994;269: 5369–5376.

14. Fernandez AM, Kim JK, Yakar S, Dupont J, Hernandez-Sanchez C, Castle AL, et al. Functional inactivation of the IGF-I and insulin receptors in skeletal muscle causes type 2 diabetes. Genes Dev. 2001;15: 1926–1934.

15. Xu E, Kumar M, Zhang Y, Ju W, Obata T, Zhang N, et al. Intra-islet insulin suppresses glucagon release via GABA-GABAA receptor system. Cell Metab. 2006;3: 47–58.

16. Halperin F, Lopez X, Manning R, Kahn CR, Kulkarni RN, Goldfine AB. Insulin augmentation of glucose-stimulated insulin secretion is impaired in insulin-resistant humans. Diabetes. 2012;61: 301–309.

17. Donath MY, Boni-Schnetzler M, Ellingsgaard H, Ehses JA. Islet inflammation impairs the pancreatic beta-cell in type 2 diabetes. Physiology (Bethesda). 2009;24: 325–331.

18. Rubino F, Forgione A, Cummings DE, Vix M, Gnuli D, Mingrone G, et al. The mechanism of diabetes control after gastrointestinal bypass surgery reveals a role of the proximal small intestine in the pathophysiology of type 2 diabetes. Ann Surg. 2006;244: 741–749.

19. Rajagopalan H, Cherrington AD, Thompson CC, Kaplan LM, Rubino F, Mingrone G, et al. Endoscopic duodenal mucosal resurfacing for the treatment of type 2 diabetes: 6-month interim analysis from the first-in-human proof-of-concept study. Diabetes Care. 2016;39: 2254–2261.

20. Kulkarni RN, Bruning JC, Winnay JN, Postic C, Magnuson MA, Kahn CR. Tissue-specific knockout of the insulin receptor in pancreatic beta cells creates an insulin secretory defect similar to that in type 2 diabetes. Cell. 1999;96: 329–339.

21. Knoop KA, McDonald KG, Kulkarni DH, Newberry RD. Antibiotics promote inflammation through the translocation of native commensal colonic bacteria. Gut. 2016;65: 1100–1109.

22. Wlodarska M, Willing B, Keeney KM, Menendez A, Bergstrom KS, Gill N, et al. Antibiotic treatment alters the colonic mucus layer and predisposes the host to exacerbated *Citrobacter rodentium*-induced colitis. Infect Immun. 2011;79: 1536–1545.

23. Kulkarni DH, McDonald KG, Knoop KA, Gustafsson JK, Kozlowski KM, Hunstad DA, et al. Goblet cell associated antigen passages are inhibited during *Salmonella typhimurium* infection to prevent pathogen dissemination and limit responses to dietary antigens. Mucosal Immunol. 2018;11: 1103–1113.

24. Paoletti AC, Parmely TJ, Tomomori-Sato C, Sato S, Zhu D, Conaway RC, et al. Quantitative proteomic analysis of distinct mammalian mediator complexes using normalized spectral abundance factors. Proc Natl Acad Sci U S A. 2006;103: 18928–18933.

