## Supplement Materials and Methods for "Linkage of a plasma zinc signature and impaired insulin receptor activation: Implications for the mechanism of type 2 diabetes mellitus"

**Supplemental Material**

***Albumin/Ig Depletion***

PureProteome™ beads (10.8 ml) were used to deplete 300 µl of plasma. The beads were separated from buffer solution using a magnet and washed once with 1x phosphate-buffered saline (PBS). Plasma was diluted 4-fold with 1x PBS to 1.2 ml, added to the washed beads and incubated for 60 min at room temperature with end-over end rotation. Following incubation, the supernatant was separated from the beads, collected and washed with 6 ml of 1x PBS three times. The collected solution was concentrated using a 3000 MWCO Amicon® Ultra-15 filter (Millipore UFC900308, Burlington MA US) at 4000*g* according to the manufacturer’s instructions. The protein concentration of the plasma was measured at 280 nm using a NanoDrop UV-Vis spectrometer (Thermo Fisher, Waltham MA US) before and after ultrafiltration to check for the integrity of the filter membrane.

***Zinc Immobilized Affinity Metal Chromatography***

Briefly, the Zn-IMAC column was washed with HIS-Select^®^ Wash Buffer (Millipore H5288, Burlington, MA US) with a volume equal to 5x of gel bed volume of 2 ml. Approximately 200 µl of sample was loaded onto the column, and flow-through was discarded. The column was eluted with HIS-Select^®^ Elution Buffer (Millipore H5413, Burlington, MA US) with a volume equal to 7x of the gel bed volume of 2 ml. The elute was concentrated using a 3000 MWCO Amicon^®^ Ultra-15 filter (Millipore UFC900308) by centrifugation at 4000*g* on a Beckman Coulter centrifuge according to the manufacturer’s instructions. Finally, the resulting concentrated sample of 240 µl was diluted 1.2-fold compared to the original unprocessed plasma.

***Liquid Chromatography with Tandem Mass Spectrometry Analysis***

At Poochon Scientific, samples provided after Zn-IMAC purification were loaded onto a peptide trap cartridge at a flow rate of 5 μL/minute. The trapped peptides eluted onto a reversed-phase 20 cm C18 PicoFrit column (New Objective, Woburn, MA) using a linear gradient of acetonitrile (3-36%) in 0.1% formic acid. The elution duration was set to 60 minutes at a flow rate of 0.3 μL/minute.

Raw data files were searched against a human protein sequence database and microbiome (bacteria) protein sequence database downloaded from the UniProt Knowledgebase (UniProtKB) protein databases website using Proteome Discoverer 2.2 software (Thermo Fisher, San Jose, CA) based on the SEQUEST and percolator algorithms. The false discovery rate (FDR) was set at 1%. The resulting Proteome Discoverer Report contains identified proteins with peptide sequences and peptide spectrum match counts (PSM#). The estimation of relative abundance for each identified protein was calculated using the normalized spectral abundance factors (NSAFs) method (24).

***Insulin Receptor Activation Assay***

The PathHunter^®^ assay uses Enzyme Fragment Complementation (EFC) technology, where the β-galactosidase (β-Gal) enzyme is split into two fragments, ProLink™ (PK) and Enzyme Acceptor (EA). Independently, these fragments have no β-Gal activity; however, when forced to complement through protein-protein interactions, they form an active β-Gal enzyme. In the PathHunter^®^ assay approach for tyrosine kinases, the ProLink tag is fused to the C-terminus of the receptor, in this case the insulin receptor. The EA is fused to a phosphotyrosine SH2 domain containing the enzyme acceptor protein that is able to bind the activated receptor tyrosine kinase. Ligand-induced activation of the receptor results in receptor phosphorylation. The SH2-EA fusion protein binds the phosphorylated receptor, forcing complementation of PK and EA to form an active β-Gal enzyme. β-Gal enzymatic activity is quantitatively measured using a chemiluminescent substrate in the PathHunter^®^ Bioassay Detection Kit.
